# XenoCell: classification of cellular barcodes in single cell experiments from xenograft samples

**DOI:** 10.1101/679183

**Authors:** Stefano Cheloni, Roman Hillje, Lucilla Luzi, Pier Giuseppe Pelicci, Elena Gatti

**Author notes:** These authors contributed equally (Stefano Cheloni, Roman Hillje). These authors contributed equally (Pier Giuseppe Pelicci, Elena Gatti).

## Abstract

Single-cell sequencing technologies provide unprecedented opportunities to deconvolve the genomic, transcriptomic or epigenomic heterogeneity of complex biological systems. Its application in samples from xenografts of patient-derived biopsies (PDX), however, is limited by the presence in the analysed samples of a mixture of cells arising from the host and the graft.

We have developed XenoCell, the first stand-alone pre-processing tool that performs fast and reliable classification of host and graft cellular barcodes. We show its application on a single cell dataset composed by human and mouse cells.

**Availability and implementation:** XenoCell is available for non-commercial use on GitLab: https://gitlab.com/XenoCell/XenoCell

## Introduction

Patient-derived xenografts (PDX) are being increasingly recognized as relevant preclinical models in many areas of biomedical research, including oncology and immunology. In recent years, the development and rapid diffusion of ultra-high-throughput droplet-based single-cell sequencing technologies has allowed resolution of genomic, transcriptomic and epigenomic profiles at the level of individual cells (Svensson, 2018). This approach proved to be invaluable for the analyses of complex and/or heterogenous biological systems, and will be increasingly used to analyse human xenograft samples (Hwang, 2018).

One potential limit of single-cell sequencing experiments of xenograft samples is the presence of host (e.g. mouse) cells along with graft (e.g. human) cells. Moreover, for reasons that are inherent to the droplet technology, a cell originating from the host may accidentally be encapsulated in the same droplet of a cell of the graft, forming a mixed-species multiplet. While several solutions have been proposed to the identification of multiplets (Wolock, 2019; McGinnis, 2019), few approaches are available to reduce host-cell contamination.

Contamination may be reduced using upstream physical or biochemical strategies such as flow cytometry-based cell sorting or laser microdissection. Downstream *in silico* techniques have been developed to separate human-mouse chimeric data by classifying individual reads, but are limited to NGS experiments on bulk cell populations (Xenome; Conway, 2012). To date, there are no tools available to pre-process chimeric data from ultra-high-throughput droplet-based single cell sequencing experiments of xenograft samples. Here, we propose XenoCell to overcome this challenge, by extending the functionality of Xenome for the classification of individual droplets and separation of cells from different organisms.

### Features and implementation

XenoCell is implemented in Python and made available as a Docker image, which contains all third-party software dependencies as a pre-configured system in order to facilitate portability across platforms and easier integration into existing workflows, such as those built with the Snakemake or Nextflow frameworks.

The XenoCell procedure consists of two main steps schematically represented in Fig 1A, starting from paired R1 and R2 FASTQ files (input) containing, respectively, barcodes and genomic cDNA sequence from droplet-based single-cell RNA or DNA sequencing experiments.

**Figure 1:**
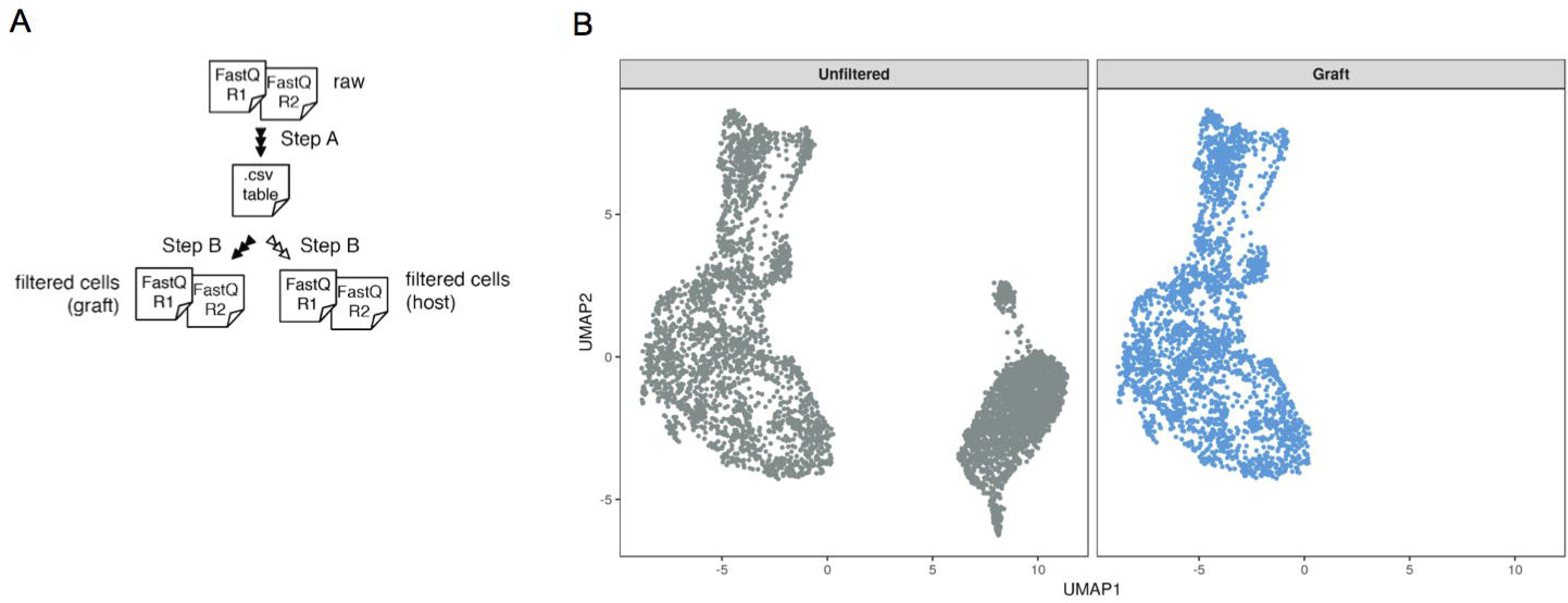
Schematic workflow of XenoCell. (A) Overview of XenoCell analysis workflow. (B) UMAP generated with unfiltered cells (gray) and graft-specific cells (blue) after alignment to the human genome (hg19). As reflected by the full overlap of the graft cells with the left group of the unfiltered cells, XenoCell successfully extracted only graft cells and, more importantly, without affecting their transcriptomic profiles.

#### Step A

XenoCell classifies each read into one of five classes (graft-specific, host-specific, ambiguous, both, and neither), retrieves the corresponding barcode-containing read for reads unambiguously assigned to either graft or host, and creates a CSV table as an intermediate output. This table contains the fractions of graft- and host-specific reads for each cellular barcode. In this step, XenoCell takes advantage of Xenome (Conway, 2012) to ensure highly accurate classification.

#### Step B

XenoCell extracts graft-specific cellular barcodes based on a user-defined upper/lower thresholds for the tolerated fraction of host/graft-specific reads (Fig. S1) (with respect to all reads associated with the respective cellular barcode).

In a separate (optional) step, users can repeat step B with a different thresholds to extract host-specific cellular barcodes, thereby allowing a separate analysis of the host cells.

The final output of XenoCell consists of filtered, paired FASTQ files which are ready to be analysed by any standard bioinformatic pipeline for single-cell analysis, such as Cell Ranger (Zheng, 2017) as well as custom workflows, e.g. based on STAR (Dobin, 2013), Seurat (Satija, 2013) and Scanpy (Wolf, 2018).

## Results and Discussion

To assess the performance of XenoCell, we used it on a publicly available single-cell gene expression dataset released by 10x Genomics (supplementary data) which is composed of a 1:1 mixture of fresh frozen human (HEK293T) and mouse (NIH3T3) cells (total of 4,952 cells).

We retrieved graft- and host-specific cellular barcodes, containing a maximum of 10% or a minimum of 90% of host-specific reads, respectively. Then, we used the XenoCell-filtered FASTQ output files as input for Cell Ranger to align the reads against the hg19 and mm10 genome, respectively, resulting in 2,532 graft and 2,433 host cells, reflecting the initial 1:1 mixture of cells.

We compared our results against the Cell Ranger pipeline, and, taking advantage of a function from Cell Ranger able to create a reference for multiple species, we aligned the human-mouse mixed dataset to a reference genome containing both human (hg19) and mouse (mm10). Cell Ranger alone resulted in slightly more cells, as compared to the filtering steps of XenoCell. Consistently, the initial 1:1 proportion of human and mouse cells was maintained. Moreover, all cells identified as graft- and host-specific by XenoCell were classified concordantly by Cell Ranger (Fig S3).

To check whether the XenoCell-filtering of cellular barcodes affects the transcriptional profiles of the single cells, we aligned the unfiltered sample and the graft-specific cells retrieved by XenoCell to the human reference genome (hg19) using Cell Ranger, and represented the transcriptional profiles in a UMAP projection generated with Seurat (Fig. 1B). Results show clearly that the graft cells retrieved by XenoCell occupy the same transcriptional space as the unfiltered sample, with the second cluster of cells likely representing mouse cells. Additionally, we aligned the unfiltered sample and host-specific cells retrieved by XenoCell to the mouse reference genome (mm10) using the same procedure as for the human cells (Fig. S2). Results lead to the same conclusion that XenoCell does not systematically alter the transcriptional profiles of the investigated cells.

Overall, XenoCell and the multi-species analysis with Cell Ranger produced successfully concordant results. The used function of Cell Ranger is available only for samples generated by the 10x Genomics kits, thereby limiting its applicability. Our tool offers instead the flexibility to set a threshold on the permitted fraction of host/graft-specific reads, depending on the biological question the user poses, and is not restricted to any library preparation kit.

## Conclusions

XenoCell is the first stand-alone tool that is able to classify and separate cellular barcodes in droplet-based single-cell (sc) sequencing experiments of xenograft samples. It has a broad range of applications, including scRNA, scDNA, scCNV, scChIP, scATAC from any combination of host and graft species. XenoCell provides paired FASTQ files as outputs, allowing large flexibility for further analysis. In conclusion, the proposed tool addresses the urgent needs of software support for analyses of single-cell data.

## Acknowledgements

Stefano Cheloni and Roman Hillje are PhD students within the European School of Molecular Medicine (SEMM).

## Supplementary Information

Supplementary data are available.

---

**Supplementary Figure 1:**
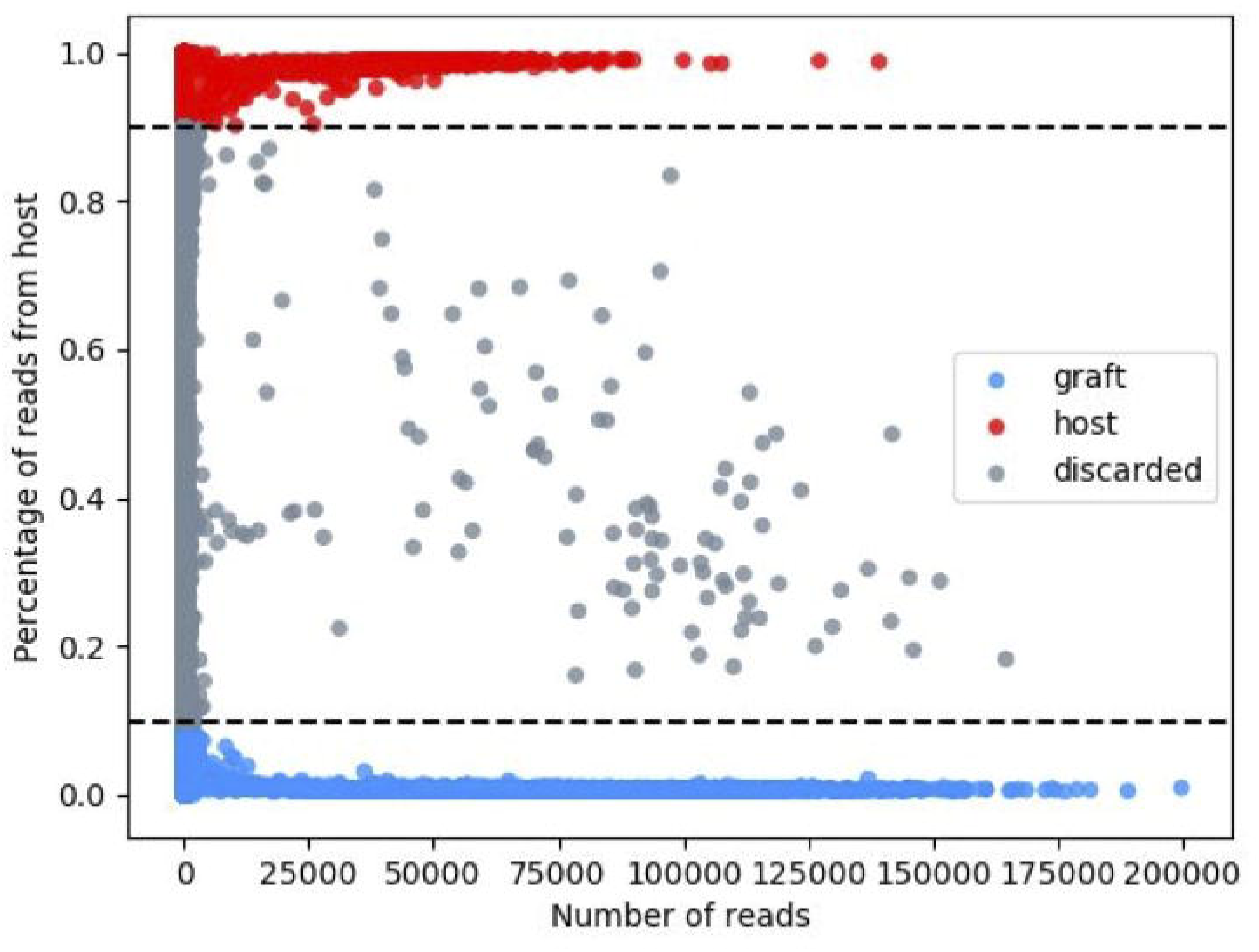
Extracting subsets of cellular barcodes based on specified thresholds. Each dot represents a cellular barcode and its corresponding percentage of reads coming from the host. Depicted dashed lines indicate the thresholds used for XenoCell (0-0.1 for graft-specific cellular barcodes; 0.9-1.0 for host-specific cellular barcodes).

**Supplementary Figure 2:**
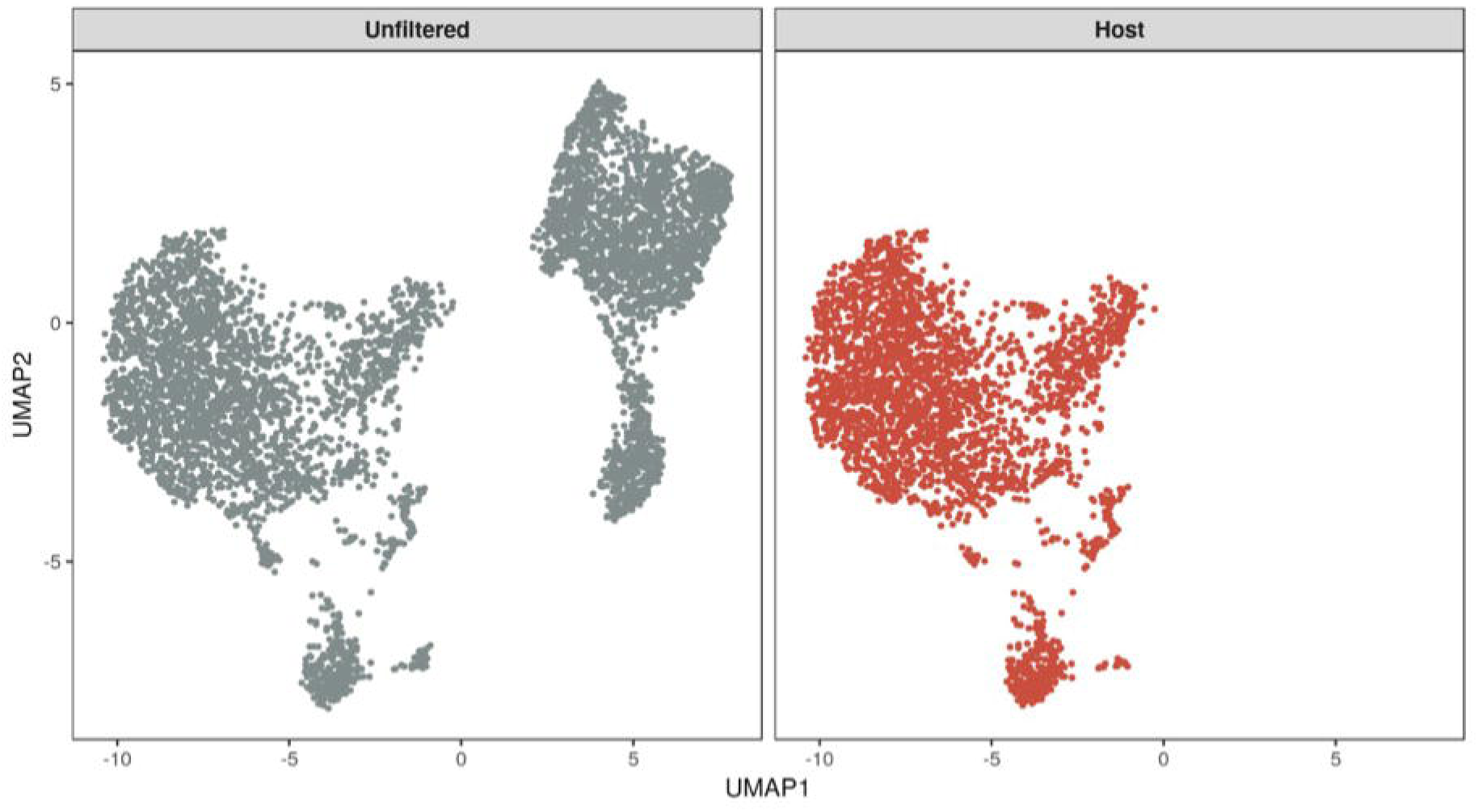
UMAP generated with unfiltered cells (gray) and host-specific cells (red) after alignment to the mouse genome (mm10). As reflected by the full overlap of the graft cells with the right group of the unfiltered cells, XenoCell successfully extracted only host cells and, more importantly, without affecting their transcriptomic profiles.

**Supplementary Figure 3:**
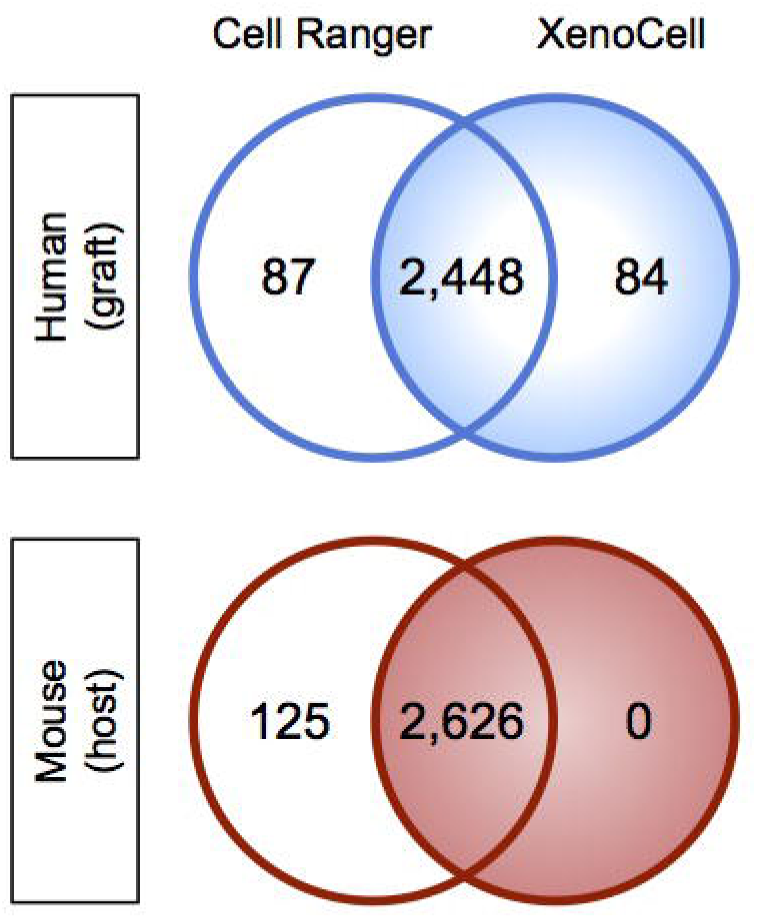
Comparison of barcode classification by XenoCell and Cell Ranger on a mixed human-mouse dataset. Both tools extract mostly the same human cells (−97% overlap), with only a few cells specific to each tool. Instead, all murine cells extracted by XenoCell were also found by Cell Ranger. The classification of cellular barcodes which were extracted by both XenoCell and Cell Ranger are concordant in all cases.

