## Supplementary Information for "XenoCell: classification of cellular barcodes in single cell experiments from xenograft samples"

The example dataset was generated using the 10x Genomics Chromium Single Cell 3’ Library (v3 chemistry) and then sequenced on Illumina NovaSeq machine. It is available through the 10x Genomics website (<https://support.10xgenomics.com/single-cell-gene-expression/datasets/3.0.0/hgmm_5k_v3> ) and licensed under the Creative Commons Attribution license.

The downstream data analysis, after XenoCell pre-processing, involved the following steps and relative tools:

- FASTQ cells were processed using Cell Ranger v3.0.2, including alignment and cell barcode filtering.
- Resulting count matrices were merged and processed in R v3.5.3 and, among other tools, Seurat v3.0.

All commands and steps, together with further information regarding the installation process and usage details, are documented and freely available through the XenoCell GitLab repository (https://gitlab.com/XenoCell/XenoCell)

We also provide a minimal Snakemake-based (Köster and Rahmann, 2012) XenoCell pipeline that allows to run the previously described steps on multiple samples in an automatised and parallelised fashion, ideally performed on high performance computing clusters due to memory requirements.

Computing time of step A for the previously described dataset was 1h 56m (16 threads), while computing time of step B (threshold 0.9) took 1h 17m on a single CPU to finish.
